# Ribosome profiling reveals ribosome stalling on tryptophan codons upon oxidative stress in fission yeast

**DOI:** 10.1101/2020.04.23.054833

**Authors:** Angela Rubio, Sanjay Ghosh, Michael Mülleder, Markus Ralser, Juan Mata

## Abstract

Modulation of translation is an essential response to stress conditions. We have investigated the translational programmes launched by the fission yeast *Schizosaccharomyces pombe* subject to five environmental stresses: oxidative stress, heavy metal, heat shock, osmotic shock and DNA damage. We also explored the contribution of two major defence pathways to these programmes: The Integrated Stress Response, which directly regulates translation initiation, and the stress-response MAPK pathway. To obtain a genome-wide and high-resolution view of this phenomenon, we performed ribosome profiling of control cells and of cells subject to each of the five stresses mentioned above, both in wild type background and in cells in which the Integrated Stress Response or the MAPK pathway were inactivated.

Translational changes were partially dependent on the integrity of both signalling pathways. Interestingly, we found that the transcription factor Fil1, a functional homologue of the Gcn4 and Atf4 proteins (from budding yeast and mammals, respectively), was highly upregulated in most stresses. Consistent with this result, Fil1 was required for the normal response to most stresses. A large group of mRNAs were translationally downregulated, including many required for ribosome biogenesis. Overall, our data suggest that severe stresses lead to the implementation of a universal translational response, which includes energy-saving measures (reduction of ribosome production) and induction of a Fil1-mediated transcriptional programme.

Surprisingly, ribosomes stalled on tryptophan codons specifically upon oxidative stress, a phenomenon that is likely caused by a decrease in charged tRNA-Tryptophan. Tryptophan stalling led to a mild translation elongation reduction and contributed to the inhibition of initiation by the Integrated Stress Response. Taken together, our results show that different stresses elicit common and specific translational responses, revealing a special and so far unknown role in Tryptophan-tRNA availability.

## INTRODUCTION

Cells react to stress situations such as starvation, changes in temperature, or the presence of toxic substances in their environment, by transcriptome and translation remodelling, as well as by the reconfiguration of their metabolism. Translation is the most energy-consuming process of the cell. Therefore, translational control plays an essential role by determining the rate of protein synthesis, which helps shape the composition of the proteome. Compared to transcriptional regulation, translational control of existing mRNAs allows for more rapid changes in protein levels, making this process particularly important upon stress exposure [1]. In addition, as protein synthesis requires a large proportion of the cell energy, its regulation is related to the metabolic status of the cell [2]. Therefore, a tight control of translation is essential to cope with stress situations, and misregulation of this process often leads to disease [3].

In eukaryotes, translational control is often performed at the initiation stage [4], when the AUG start codon is identified and decoded by the initiator tRNA (Met-tRNAi Met). A key regulator of this process is the translation initiation factor eIF2, which is part of the so-called ternary complex (TC) together with the initiator tRNA and GTP. The TC, in complex with other translation initiation factors, binds to the 40S ribosomal subunit to form the 43S preinitiation complex (PIC). The 43S PIC is recruited to the 5’ cap of the mRNA by additional initiation factors, leading to the formation of the 48S PIC, which scans the mRNA until the initiation codon is reached. At this step, the 60S subunit binds the complex and GTP is hydrolysed. eIF2-GDP must then be recycled to eIF2-GTP by the GTP/GDP-exchange factor eIF2B. After stress exposure, the eIF2α subunit is phosphorylated at a specific serine residue (serine 51 in mammals and 52 in fission yeast), and binds to eIF2B acting as a competitive inhibitor, which reduces levels of the ternary complex and triggers a global down-regulation of translation [5,6]. The pathway that regulates translation through eIF2α phosphorylation is called the Integrated Stress Response. In mammals, there are four eIF2α kinases (Hri, Gcn2, Pek/Perk and Pkr) that are activated by different stresses and inhibit translation initiation through the phosphorylation of eIF2α [4]. In the yeast *Saccharomyces cerevisiae*, Gcn2 is the sole eIF2α kinase [7,8]. In the fission yeast *Schizosaccharomyces pombe*, three eIF2α kinases (Gcn2, Hri1 and Hri2) show distinct and overlapping activation patterns in response to cellular stresses [9,10]. Whereas Hri2 is mainly activated in response to heat shock and Hri1 at stationary phase in response to nutritional limitation, Gcn2 is the main eIF2α kinase activated in early exposure to H_2_O_2_ and MMS [9–11].

In parallel to the general downregulation of translation upon stress, there is an induction of the translation of specific mRNAs, some of them encoding transcription factors, which in turn promote the transcriptional response. Recently, we have shown that amino acid starvation in *S. pombe* increases the translation of the transcription factor Fil1, the functional orthologue of Atf4 in mammals and Gcn4 in budding yeast, through the activation of the Gcn2-eIF2α pathway [12]. Fil1 is required for the transcriptional response to amino acid starvation as well as for normal growth in minimal medium lacking amino acids. Furthermore, Fil1 is regulated in a similar manner through inhibitory upstream ORFs (uORFs) located at the 5’-leader sequence (six uORFs in *fil1*, four in *GCN4* and two in *ATF4*) [12]. In budding yeast and mammalian cells, eIF2α phosphorylation reduces the abundance of active ternary complexes and the reinitiation of translation occurs after bypassing the inhibitory uORFs, which allows the scanning subunit to reach the main coding sequence [8,13]. Notably, Fil1 does not show sequence similarity to either Gcn4 or Atf4 [12].

Gene expression programs in response to stress are also regulated by Stress Activated Protein Kinases (SAPK). A key player in this pathway in *S. pombe* is the mitogen-activated protein kinase (MAPK) Sty1/Spc1 [14,15], which is homologous to the Hog1 osmo-sensing MAPK in *S. cerevisiae* and to the mammalian and *Drosophila* JNK and p38 SAPKs [16].

Stress signals activate and phosphorylate Sty1, promoting its transient accumulation in the nucleus, where it triggers a wide transcriptional shift of the gene expression program. This transcriptional response is mediated by the Atf1 transcription factor [17–20]. Sty1 also has a role in the translational response to stress. First, cells show higher levels of eIF2α phosphorylation in the absence of Sty1 [10,21]. Second, Sty1 associates *in vivo* with the translation elongation factor 2 (eEF2) and the translation initiation factor 3a (eIF3a) [22]. Finally, the presence of Sty1 is required to maintain the levels and the phosphorylation of eIF3a, and the recovery of translation levels after stress is less efficient in the absence of Sty1 [21,22].

Translation upon stress can be also regulated at the elongation step through the abundance, modification, and charging levels of transfer RNA (tRNA) [23]. tRNA is the most extensively modified RNA and many post-transcriptional nucleoside modifications occur at the anticodon loop [24]. These modifications can change the stability or localization of tRNAs [25], as well as the fidelity and efficiency of translation [26]. It has also been shown that tRNA fragmentation occurs upon stress in eukaryotes [27] and, in human cells, that tRNA fragments have a role in translation repression [28].

Ribosome profiling (ribo-seq) provides a genome-wide and high-resolution view of translational control. The approach is based on the treatment of translating ribosome–mRNA complexes with a ribonuclease (RNase), in such a way that only RNA fragments protected by bound ribosome survive the treatment. These fragments are then isolated and analysed by high-throughput sequencing. The number of sequence reads that map to a coding sequence, normalized by mRNA levels, provides an estimate of the efficiency of translation for every cellular mRNA [29]. In addition, the position of the reads on the genome identifies the location of ribosomes on the mRNA. This information can be used to determine relative ribosome occupancies on each codon, which allow the detection of ribosomes stalled on specific codons [12,30,31].

Although some stress-induced gene expression programs have been studied in detail at the genome-wide level [14,15,32–36], the effects of different stresses have been examined in isolation, making comparisons across stresses difficult. Moreover, to our knowledge, there are no systematic studies of the role of major stress-response pathways on genome-wide translation programmes. Here we use *S. pombe* to investigate similarities and differences among the translational responses to five commonly studied stress situations: exposure to H_2_O_2_ (oxidative stress), cadmium (heavy metal), and methyl methanesulfonate (MMS, genotoxic stress), sorbitol treatment to induce osmotic shock, and heat shock. We perform ribosome profiling in control and stressed cells, under highly controlled and comparable conditions. We also investigate the contribution of the Integrated Stress Response (mediated by eIF2α phosphorylation) and the stress-responsive MAPK pathway (Sty1) to these translation programs. Overall, we found that translation is typically downregulated, but activated for very few genes. Interestingly, the Fil1 transcription factor is strongly induced at the translational level upon several stresses, in a manner strongly dependent on eIF2α phosphorylation. Moreover, Fil1 is required for the full implementation of stress-responsive transcription programs. These stresses also cause a rapid translational downregulation of genes involved in ribosome biogenesis and ribosomal proteins.

Surprisingly, we found that ribosomes stall selectively on tryptophan codons upon oxidative stress, a phenomenon that is likely to be caused by a decrease in the levels of charged tRNA-Tryptophan (tRNA-Trp). This specific stalling on tryptophan led to a mild elongation defect. Our results suggest that different stresses elicit common and specific translational responses, both at the initiation and the elongation levels, and uncovers a novel and specific role to tryptophan tRNA availability.

## RESULTS

### Transcriptomic responses to stress

To investigate genome-wide effects of stress on gene expression, we carried out two independent ribosome profiling experiments (with parallel RNA-seq) of *S. pombe* cells subject to five different stress-inducing treatments for 15 minutes: 0.5 mM H_2_O_2_ (oxidative stress), 0.5 mM CdSO_4_ (heavy metal exposure), temperature shift from 32°C to 39°C (heat shock), 1 M sorbitol (osmotic shock) and 0.02% methyl methanesulfonate (MMS, an alkylating agent that causes DNA damage). To study the contribution of the eIF2α phosphorylation and Sty1 pathways we performed parallel experiments with the mutant strain *eIF2α-S52A*, which expresses a non-phosphorylatable version of eIF2α, and a *sty1Δ* strain, in which the main stress-responsive MAPK pathway is inactive. These conditions have been used in the past for microarray-based transcriptomics, and have been shown to elicit robust transcriptional responses within a similar timeframe while maintaining high cell viability [14]. All experiments were carried out using rich medium (YE), as *sty1Δ* cells grow very poorly in minimal medium. We first examined the transcriptional responses to stress using the RNA-seq data. As we did not perform a global correction for total mRNA abundance, the discussion below refers to relative changes in gene expression. There were more genes significantly up-regulated than down-regulated in every strain and condition (Table 1). For the specific stress conditions that we employed (e.g. concentration of stressor and time), the changes were stronger after heat and cadmium stress, but genes induced by the five stress treatments overlapped significantly with one another. A similar overlap across stresses was observed for repressed genes (highest P value < 2× 10^−5^ in wild type cells). Figure S1A shows a heat map with over 1,200 genes that are differentially expressed in at least one condition in wild-type cells. Most of the regulated genes behave similarly in wild type and *eIF2α-S52A* cells. By contrast, the transcriptional response to stress in *sty1Δ* cells was substantially weaker although not completely abolished [14] (Fig. S1A). Consistently, the numbers of induced and repressed genes were comparable among wild type and the *eIF2α-S52A* mutant but were lower in *sty1Δ* cells (Table 1).

**Table 1.**
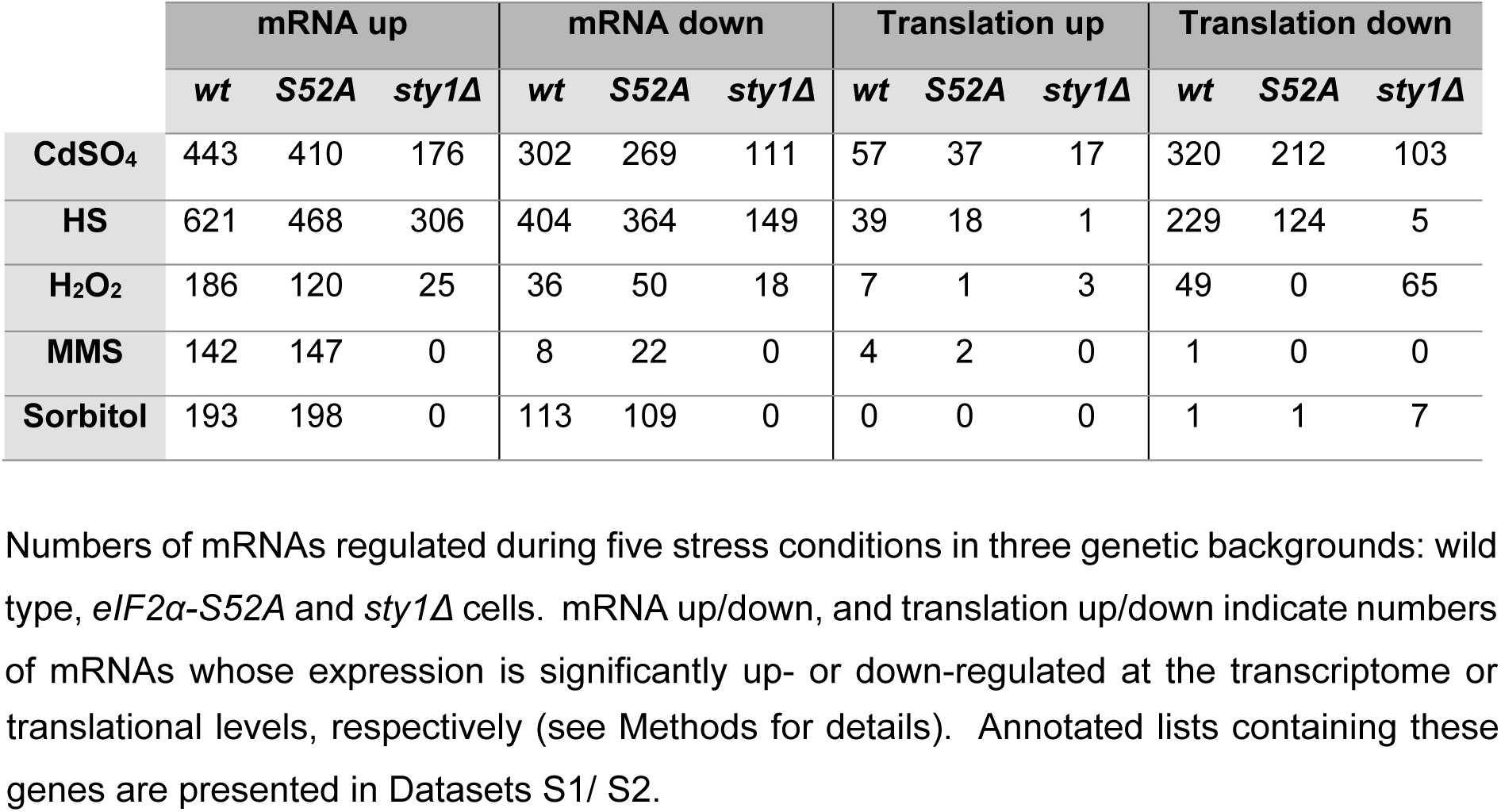
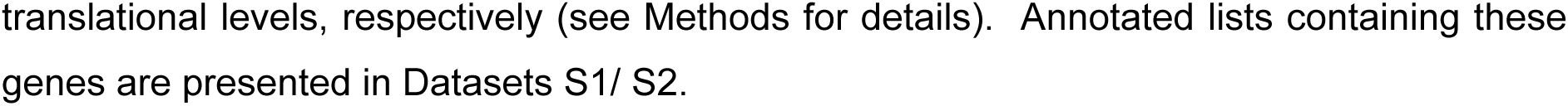
Number of transcripts regulated after stress exposure on each strain.

|  | mRNA up |  |  | mRNA down |  |  | Translation up |  |  | Translation down |  |  |
| --- | --- | --- | --- | --- | --- | --- | --- | --- | --- | --- | --- | --- |
| | *wt* | *S52A* | *sty1*Δ | *wt* | *S52A* | *sty1*Δ | *wt* | *S52A* | *sty1*Δ | *wt* | *S52A* | *sty1*Δ |
| **CdSO<sub>4</sub>** | 443 | 410 | 176 | 302 | 269 | 111 | 57 | 37 | 17 | 320 | 212 | 103 |
| **HS** | 621 | 468 | 306 | 404 | 364 | 149 | 39 | 18 | 1 | 229 | 124 | 5 |
| **H<sub>2</sub>O<sub>2</sub>** | 186 | 120 | 25 | 36 | 50 | 18 | 7 | 1 | 3 | 49 | 0 | 65 |
| **MMS** | 142 | 147 | 0 | 8 | 22 | 0 | 4 | 2 | 0 | 1 | 0 | 0 |
| **Sorbitol** | 193 | 198 | 0 | 113 | 109 | 0 | 0 | 0 | 0 | 1 | 1 | 7 |

Numbers of mRNAs regulated during five stress conditions in three genetic backgrounds: wild type, *eIF2 α-S52A* and *sty1* $\Delta$ cells. mRNA up/down, and translation up/down indicate numbers of mRNAs whose expression is significantly up- or down-regulated at the transcriptome or
translational levels, respectively (see Methods for details). Annotated lists containing these genes are presented in Datasets S1/ S2.

The Core Environmental Stress Response (CESR) was defined in a microarray study as those genes induced or repressed two-fold or greater in most of the five stresses analysed [14]. Consistent with the microarray data, the majority of induced and repressed CESR genes, as well as stress-specific genes, were also differentially expressed in our RNA-seq experiments (Fig. 1A, B and Fig. S1B, S1C). Overall, our data show that transcriptomic responses to stress are largely independent of eIF2α phosphorylation, but strongly reliant on the MAPK pathway.

**Fig. 1.**
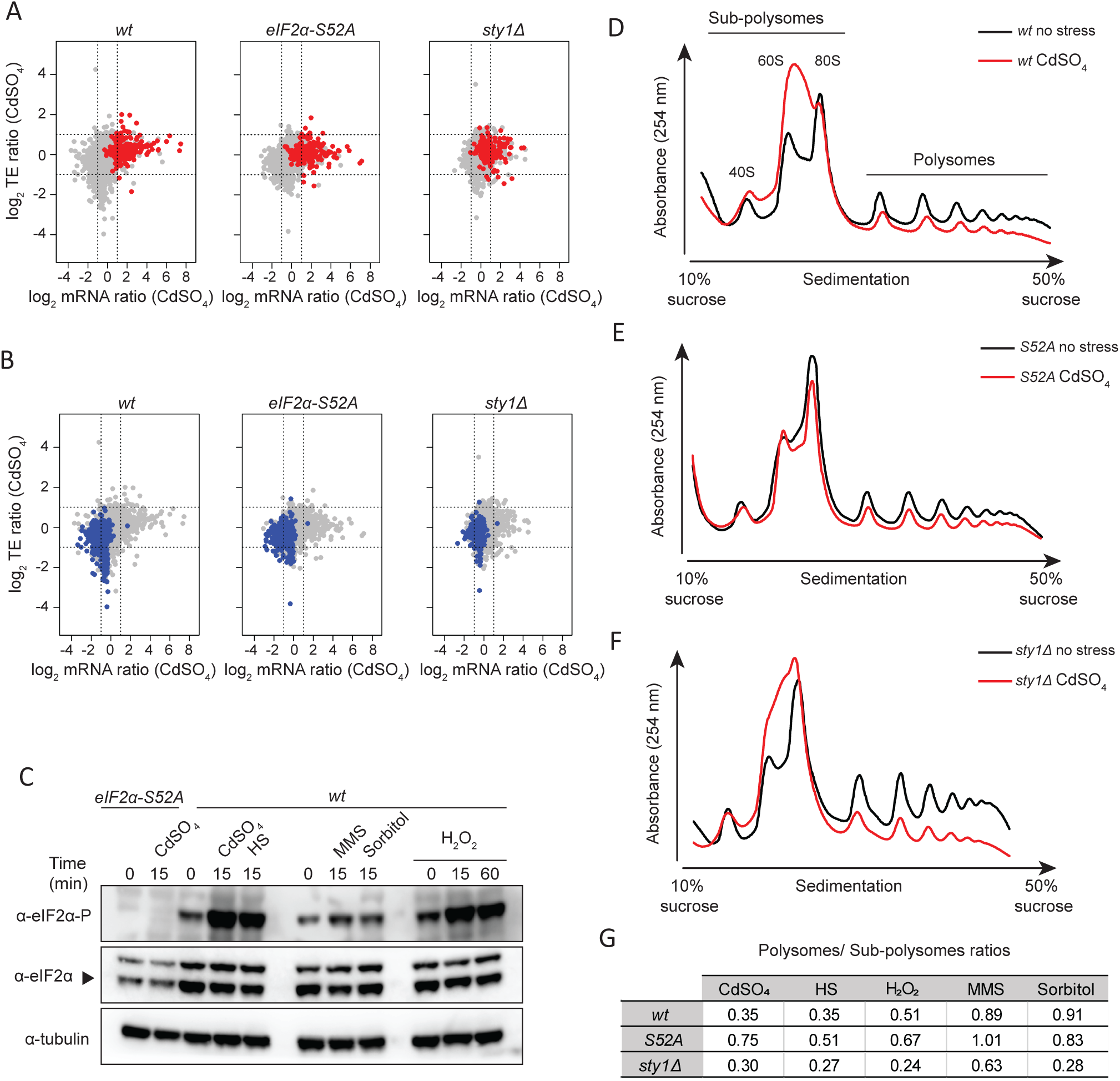
General responses to stress. **(A)** Scatter plot comparing mRNA levels and translation efficiencies (log_2_ ratios stress/control) upon cadmium treatment (15 min) in wild type, eIF2α-S52A and sty1Δ genetic backgrounds. The results of a single experiment are shown. CESR-induced genes are plotted in red. **(B)** As in A, but CESR-repressed genes are plotted in blue. **(C)** Western blots comparing eIF2α phosphorylation levels after the five treatments for the indicated times in wild type cells. In the first two lanes (left), eIF2α-S52A cells were used as negative control for the anti-eIF2α phosphorylation antibody. Tubulin was employed as a loading control. **(D to F)** Representative polysome profile traces before and after cadmium treatment (15 min) in wild type **(D)**, eIF2α-S52A **(E)** and sty1Δ cells **(F). (G)** Estimation of polysome/subpolysome ratios after five stress treatments (15 min) relative to unstressed conditions.

### General translational responses to stress

Activation of the Integrated Stress Response by stress leads to general translational down-regulation, mediated through eIF2α phosphorylation. To investigate the level of activation of this pathway and its effect on global translation levels, we monitored eIF2α phosphorylation levels by immunoblotting and analysis of polysome profiles on sucrose gradients. For the specific conditions we used (single stressor concentration and 15-minute exposures), cadmium, heat shock and H_2_O_2_ treatments led to increased eIF2α phosphorylation levels (Fig. 1C) and lower polysomes to subpolysomes ratios (Fig. 1D-G), the latter being indicative of a decrease in translation initiation. Both effects were also detected, although less pronounced, after the MMS and sorbitol treatments (Fig. 1C, 1G). *eIF2α-S52A* cells reduced the polysomes to subpolysomes ratio in response to stress, albeit to a lesser extent than wild-type cells (Fig. 1D-G). This partial dependency on eIF2α phosphorylation is in in agreement with recent reports of *S. pombe* responses to UV and oxidative stress [35,37,38]. By contrast, the Sty1 protein was not required for the downregulation of translation initiation (Fig. 1F, G). Indeed, in some cases (H_2_O_2_ and sorbitol), *sty1Δ* cells reduced translation more than wild type cells (Fig. 1G). Thus, since polysome profiling cannot distinguish between unaltered active translation and slowed-down elongation, our polysome profiling data suggest that translation is affected at least at the initiation step upon cadmium, heat shock and H_2_O_2_ exposure. This effect is only partially dependent on eIF2α phosphorylation. In addition, the Sty1 MAPK pathway may also modulate translation initiation under some stresses [10,21].

### Gene-specific translational regulation upon stress exposure

We used ribosome profiling to investigate gene-specific translational control in response to stress. Ribosome-protected fragments (RPFs) were isolated and analysed by high-throughput sequencing, whereas mRNA levels were estimated in parallel by RNA-seq. The number of RPFs mapped to each coding gene, normalized by the corresponding number of RNA-seq reads, was used to calculate relative translation efficiencies (TEs) (Fig. 2A). Genes showing significant changes in TE upon stress treatment were identified using RiboDiff (see Methods), with thresholds of a minimum 1.5-fold change and adjusted P value < 0.01. This approach does not consider global changes in gene expression and, therefore, may overestimate TE values. Nevertheless, these TE values reflect relative changes among conditions and among genetic backgrounds, and identifies genes that behave differently from the majority of transcripts.

**Fig. 2.**
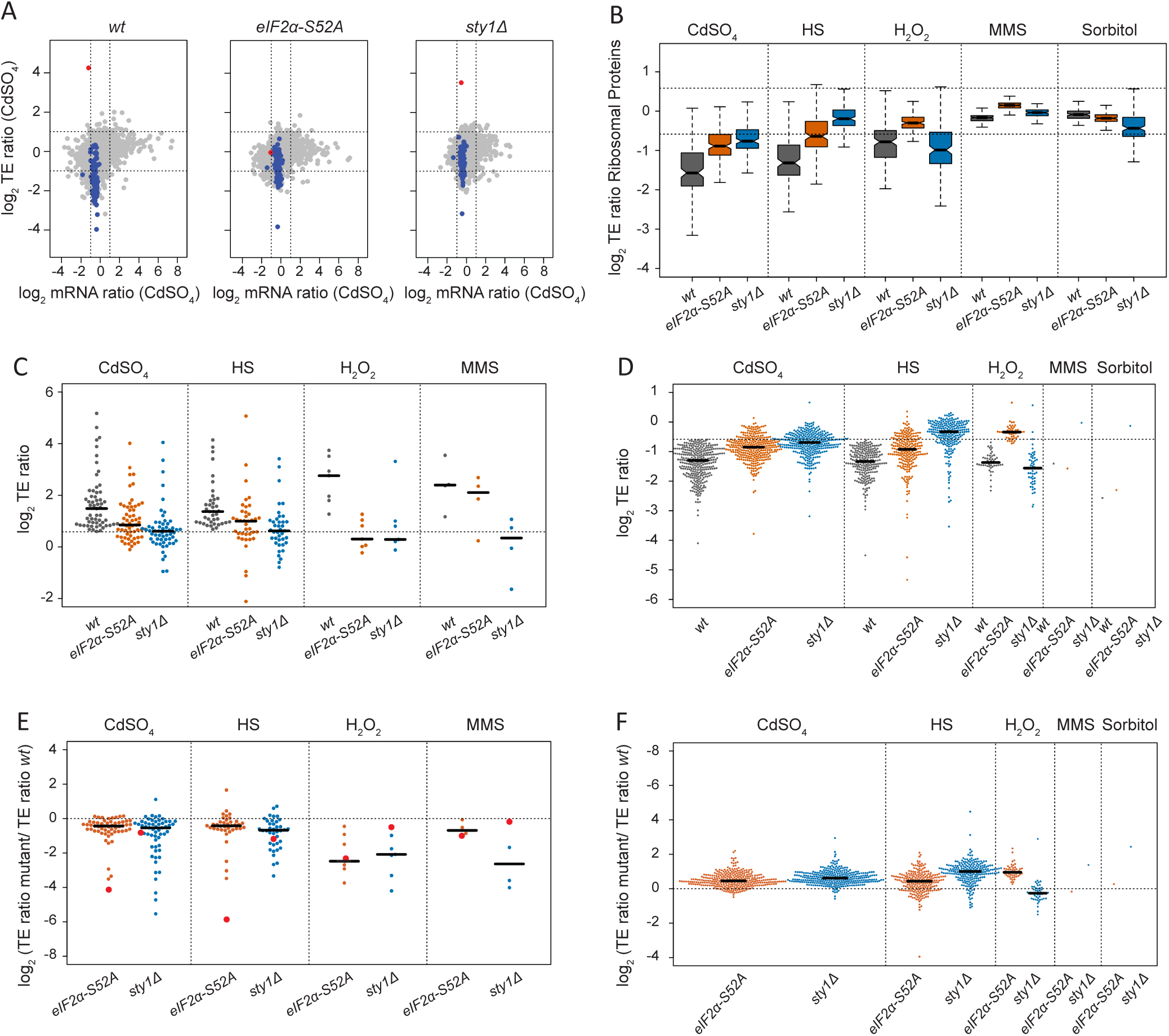
Translational regulation upon stress exposure. **(A)** Scatter plot comparing mRNA levels and translation efficiencies (log_2_ ratios stress/control) upon cadmium treatment (15 min) in wild type, *eIF2 α-S52A* and *sty1Δ* genetic backgrounds. The results of a single experiment are shown. The *fil1* gene is plotted in red and genes encoding ribosomal proteins in blue. **(B)** Boxplots comparing translation efficiencies of stressed and control cells (log_2_ ratios stress/control) of genes encoding ribosomal proteins. Data are shown for wild type, *eIF2α-S52A* and *sty1Δ* cells. **(C)** Comparisons of translation efficiencies of stressed (15 min) and control cells (log_2_ ratios stress/control). Only genes that showed significant translational upregulation in wild type cells in at least one stress are displayed. Data are presented for wild type, *eIF2α-S52A* and *sty1Δ* cells. **(D)** As in C, but only genes that showed significant translational downregulation are displayed. **(E)** As in C, but TE changes have been normalised to those of wild type cells. Dots corresponding to *fil1* gene are shown in red. **(F)** As in E, but data are displayed for significantly downregulated genes.

Numbers of mRNAs regulated during five stress conditions in three genetic backgrounds: wild type, *eIF2α-S52A* and *sty1Δ* cells. mRNA up/down, and translation up/down indicate numbers of mRNAs whose expression is significantly up- or down-regulated at the transcriptome or translational levels, respectively (see Methods for details). Annotated lists containing these genes are presented in Datasets S1/ S2.

Upon stress, the number of translationally regulated genes was much smaller than those affected at the transcript level (Table 1). In relative terms, unlike transcriptomic responses, the numbers of translationally down-regulated genes were higher than those up-regulated (Table 1). Thus, the gene-specific translational response seems to be targeted to reduce mRNA translation. The treatment that had the strongest effect was cadmium, followed by heat shock and H_2_O_2_, while fewer changes were detected upon the applied MMS and sorbitol concentrations (Table 1). This trend is generally similar to transcriptional and eIF2α phosphorylation changes (Fig. 1), indicating that gene expression programs in response to the latter stresses are generally weaker.

We found 82 translationally upregulated genes in wild type cells in at least one stress treatment, 28 of which overlapped with the CESR induced genes (P value < 5× 10^−15^). Of those 82 genes, four were induced in all the stress conditions except for sorbitol, and 15 were shared across cadmium and heat shock. We could not find any specific GO category enriched within the 82 genes (see Methods). Yet, the data included individual genes that have previously linked to the stress response. An example of cadmium-upregulated gene (both transcriptionally and translationally) was *prr1*, which encodes a transcription factor involved in the oxidative stress response and sexual differentiation [39–41] (Fig. S2A). This regulation is consistent with the generation of reactive oxygen species after cadmium stress [42]. Other interesting genes were related to protein catabolism (*ubp3*, in cadmium and heat shock), autophagy (*atg3*, in cadmium), cell cycle (*srk1*, in cadmium), DNA repair (*dna2*, in heat shock and *uve1*, in H_2_O_2_ and cadmium), transmembrane transport, carbohydrate and amino acid metabolism. The most translationally upregulated gene was *fil1* [12], which encodes a transcription factor essential for the response to amino acid starvation (Fig. 2A). The role of this gene in stress responses is discussed in detail below. Finally, we found that upon cadmium and heat shock treatments, more than 45% of translationally upregulated genes were also transcriptionally induced, whereas less than 14% of translationally downregulated were also transcriptionally repressed. This coordination of transcriptomic and translational induction (potentiation) has been observed by polysome profiling in budding and fission yeast responses to stress [36,43].

We also identified 382 translationally down-regulated genes (translation efficiency repressed in at least one condition), 149 of which were part of the CESR-repressed list (P value < 5× 10^−73^). Like the induced genes, cadmium, heat shock and H_2_O_2_ led to stronger effects, whereas the consequences of MMS and sorbitol exposure were very mild (Table 1). The most extensive overlap was between cadmium and heat shock stress, with 170 genes shared. These genes included *cdr2*, which is involved in the regulation of the G2/M transition through the inhibition of the Wee1 kinase [44] (Fig. S2B). Repression of this gene is consistent with the block of cell cycle progression in response to stress. There was also a substantial overlap (38 genes) among cadmium, heat shock and H_2_O_2_ stresses. These data indicate that heavy metal exposure, oxidative conditions and heat shock are strong stresses under the conditions used in this work, and that they cause similar translational responses.

Translationally repressed genes were mainly associated with cytoplasmic translation (GO:0002181; P = 10^−89^), ribosome biogenesis (GO:0042254; P = 10^−28^) and ribosome assembly (GO:0042255; P = 10^−11^). Consistently, 110 of the 382 down-regulated genes encoded ribosomal proteins. (Fig. 2A, B). We have previously observed this effect upon nitrogen depletion [30] and amino acid starvation [12], demonstrating that this is a widespread translational response to stress.

Finally, we investigated whether translational responses were dependent on the major stress response pathways. Many TE-upregulated genes were partially induced in both *eIF2α-S52A* and *sty1Δ* mutants (Fig. 2C). However, a direct comparison of the induction levels in wild type and mutant cells (Fig. 2E), revealed that the upregulation in the mutant was impaired for most genes. A very similar dependency was observed for downregulated genes (Fig. 2D, F), which were generally less repressed in both mutant backgrounds. Therefore, both the eIF2α and the MAPK pathways contribute to a normal translational response, but neither of them is sufficient to explain the response in its entirety.

### *Fil1* is the major translational responder to stress

The transcription factor *fil1* was induced very strongly at the translation efficiency level by cadmium, heat shock, and H_2_O_2_ treatments, while showing small reductions or no changes in mRNA levels (Fig. 3A, B). There was also a weak translational induction after MMS treatment, but none after the sorbitol doses applied. These data mirror the change in eIF2α phosphorylation in these stresses (Fig. 1C). Consistently, the translational induction of *fil1* was completely dependent on eIF2α phosphorylation, and very weakly on Sty1 signalling (Fig. 3A). We investigated if this increase in TE was accompanied by higher protein levels. To do this, cells expressing Fil1-TAP from their endogenous locus were used to compare protein levels by immunoblot in the five stress conditions. Consistent with the TE data, there was a strong increase in protein levels after cadmium, heat shock and H_2_O_2_ treatments, weaker after MMS, and none upon sorbitol (Fig. 3C, D and Fig. S2 C-E). The kinetics of the *fil1* induction was stress-specific, with cadmium and heat shock showing a transient response (peaking at 15 minutes) and H_2_O_2_ and MMS showing slower responses. In particular, the increase at 60 minutes after H_2_O_2_ and MMS (Fig. 3D and Fig. S2D) was analogous to the behaviour of CESR-induced genes under the same conditions, whose induction persisted for an hour [14]. The variety in induction kinetics suggests that Fil1 protein induction may be regulated at other levels in addition to translation. Consistent with the ribosome profiling data, no Fil1 protein induction was detected in *eIF2α-S52A* cells (Fig. 3C, D and Fig. S2C-E). Surprisingly, Fil1 protein levels were higher in *sty1Δ* than in wild type cells, even in the absence of stress (Fig. 3C, D and Fig. S2C-E). This may relate to the fact that *sty1Δ* cells are sensitive to stress [45–48]. Indeed cells lacking this protein show evidence of an induced stress response even under normal laboratory growth conditions [14]. Taken together, these data indicate that *fil1* translational induction upon stress leads to an increase in Fil1 protein levels in an eIF2α-dependent manner, and that the Sty1 MAPK pathway modulates this effect. This response takes place in rich medium, where Fil1 is not required for normal growth [12]. Therefore, Fil1 behaves as a general stress-responsive transcription factor.

**Fig. 3.**
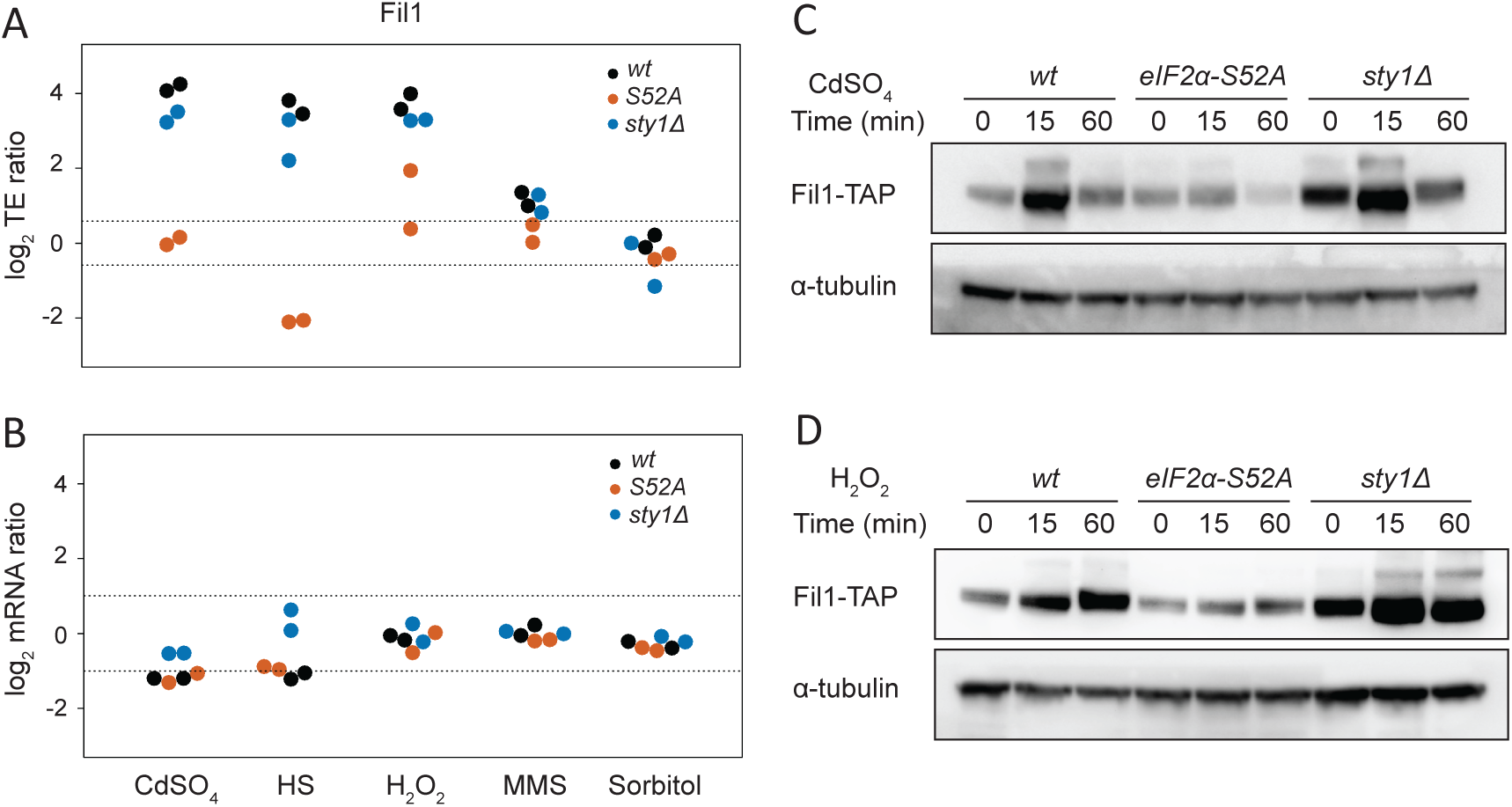
*Fil1* is the major translational responder to stress. **(A)** Comparison of translation efficiency of the *fil1* gene between stressed and control cells (log_2_ ratios stress/control). The dotted lines indicate 1.5-fold changes. Data are presented for wild type, *eIF2α-S52A* and *sty1Δ* cells. **(B)** As in C, but for *fil1* mRNA changes. The dotted lines indicate 2-fold changes **(C)** Western blots to measure Fil1-TAP protein levels after cadmium treatment for the indicated times. Data are presented for wild type, *eIF2 α-S52A* and *sty1Δ* cells. Tubulin was used as a loading control. **(D)** As in C, but after H_2_O_2_ treatment.

We then investigated the role of Fil1 in the transcriptional responses to stress. We performed RNA-seq experiments in the five stress conditions in cells lacking *fil1*, and monitored the behaviour of 165 previously identified Fil1 targets [12]. Note that Fil1 targets were defined as genes that showed lower expression in *fil1Δ* cells in minimal medium and in the absence of stress [12]. In unstressed cells growing in rich medium, Fil1 targets were expressed at slightly lower levels in the *fil1Δ* mutant (Fig. 4A-C and Fig. S3A, B). In response to cadmium, heat shock and H_2_O_2_ treatments, Fil1 targets were expressed at substantially decreased levels in the mutant (Fig. 4A-C and Fig. S3A, B). Consistently, Fil1 target genes overlapped significantly with genes underexpressed in *fil1Δ* cells at 15 minutes after heat shock (Fig. 4D), and with genes underexpressed 60 minutes after H_2_O_2_ treatment (Fig. 4E). These data demonstrate that Fil1 promotes the expression of a common group of genes in response to strong stresses (cadmium, heat shock and H_2_O_2_ treatments).

**Fig. 4.**
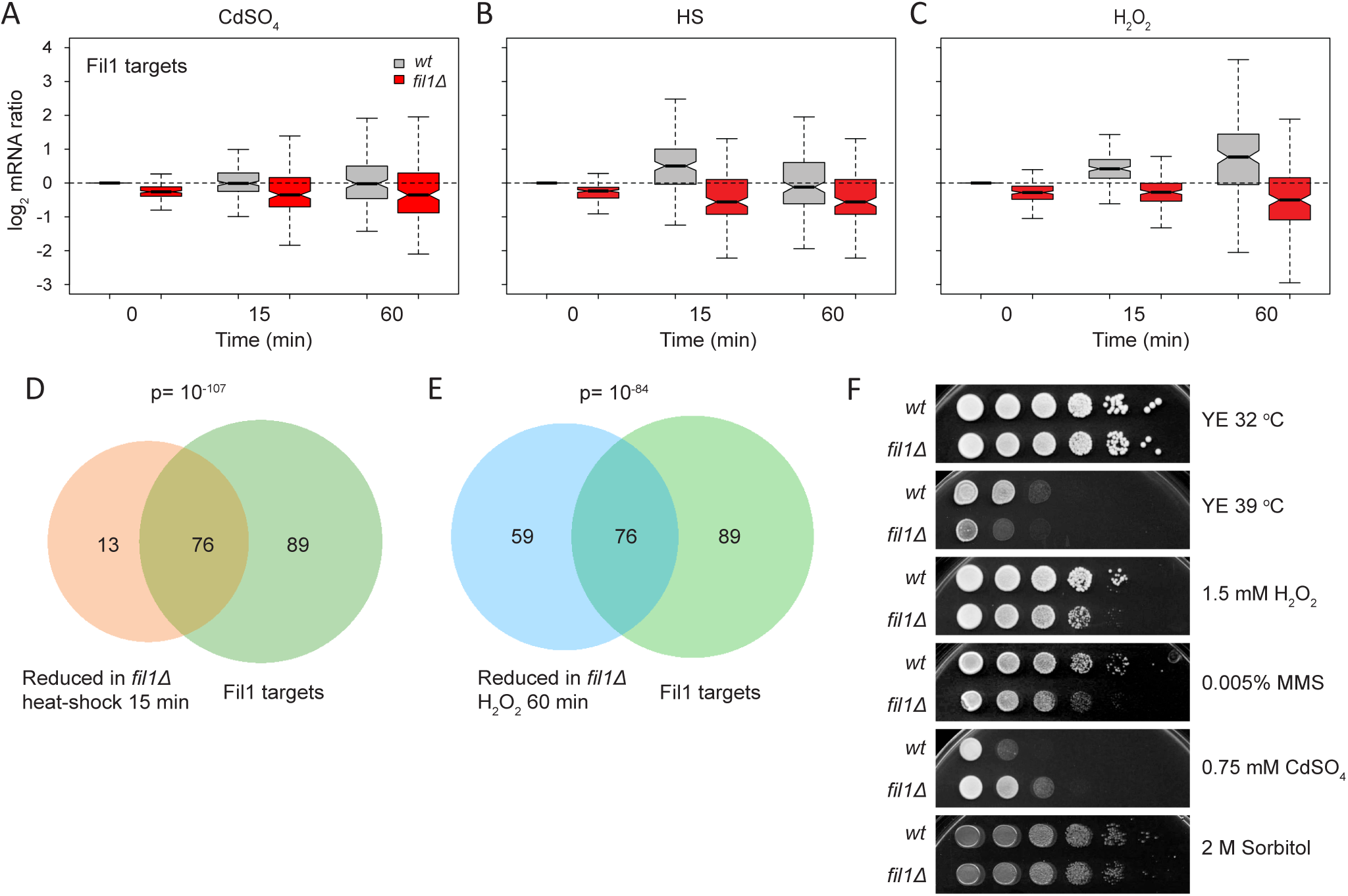
Role of Fil1 in the transcriptional responses to stress. **(A to C)** Boxplots comparing mRNA levels of stressed and control cells (log_2_ ratios stress/control) of Fil1 targets. Data are shown for wild type and *fil1Δ* cells at the indicated times and stresses **(D)** Venn diagram showing the overlap between genes expressed at low levels in *fil1Δ* mutant relative to wild type cells after heat shock (15 min), and Fil1 targets in unstressed cells. The P value of the observed overlap is shown. **(E)** As in D, but genes expressed at low levels in *fil1Δ* mutant relative to wild type after H_2_O_2_ treatment (60 min) were compared to Fil1 targets in unstressed cells. **(F)** Drop assays of wild type and *fil1Δ* cells plated in the indicated conditions.

These results suggest that Fil1 may be important for survival to stress in rich medium. We explored this hypothesis by performing viability assays of wild type and *fil1Δ* mutant under the five stress treatments. *fil1Δ* cells were sensitive to high temperature, H_2_O_2_ and MMS, whereas no difference to wild-type was observed at the sorbitol concentrations applied. Surprisingly, cells lacking Fil1 were resistant to cadmium treatment (Fig. 4F). Although the reason for this phenotype is unclear, deletion of genes encoding other transcription factors involved in stress response (*atf1*) [39,49], and of genes encoding several RNA-binding proteins [50] show similar resistance to cadmium.

As mentioned above, Fil1 is necessary for the normal expression of its targets in response to several stresses (Fig. 4A-C), and Fil1 expression was induced under the same conditions in an eIF2α-dependent manner (Fig. 3 C, D and Fig. S2C). To investigate if *fil1* induction is required for the normal expression of Fil1 targets, we compared the expression levels of *fil1* targets upon stress in wild type and eIF2α-S52A mutants. After 15 minutes of treatment, Fil1 target levels were mildly increased by heat shock and H_2_O_2_ (but not by cadmium) in an eIF2α-dependent manner (Fig. S3C). As Fil1 targets tend to be induced more strongly by H_2_O_2_ at later time points (Fig. 4C), we repeated the experiment upon a 60-minute H_2_O_2_ exposure. Indeed, this led to a late and stronger induction of Fil1 targets that was almost completely dependent on eIF2α phosphorylation (Fig. S3D). These results indicate that, at least in some conditions, *fil1* translational upregulation plays a role in the implementation of the normal transcriptional responses to stress.

In addition, cells lacking Sty1 showed increased mRNA levels of Fil1 targets in both stressed and unstressed cells (Fig. S3E). Indeed, the overlap between upregulated genes in unstressed *sty1Δ* cells and Fil1-dependent genes was significant (P= 3.5 × 10^−11^). These data are also consistent with the increased Fil1 protein levels in unstressed *sty1Δ* cells (Fig. 3C-D and Fig. S2C-E). These data suggest that the Sty1 MAPK pathway cross-talks with the Integrated Stress Response, consistent with previous observations [10,21].

### Ribosome occupancy on tryptophan codons is increased upon oxidative stress

Ribosome profiling provides information on ribosome locations with codon-level resolution and can thus be used to detect codon-specific ribosome stalling caused by stress conditions. To investigate this level of regulation, we quantified the fraction of ribosomes translating each of the 61 amino acid-encoding codons, normalized by the abundance of the corresponding codon in the transcriptome. This ‘relative codon occupancy’ reflects the average time spent by the ribosome on each codon. Strikingly, the single codon for tryptophan (TGG) showed strongly increased ribosome occupancy upon H_2_O_2_ treatment (Fig. 5A, S4A). This enrichment was highly specific, as it was not observed for any other codon, and in any other stress condition (Fig. S4B).

**Fig. 5.**
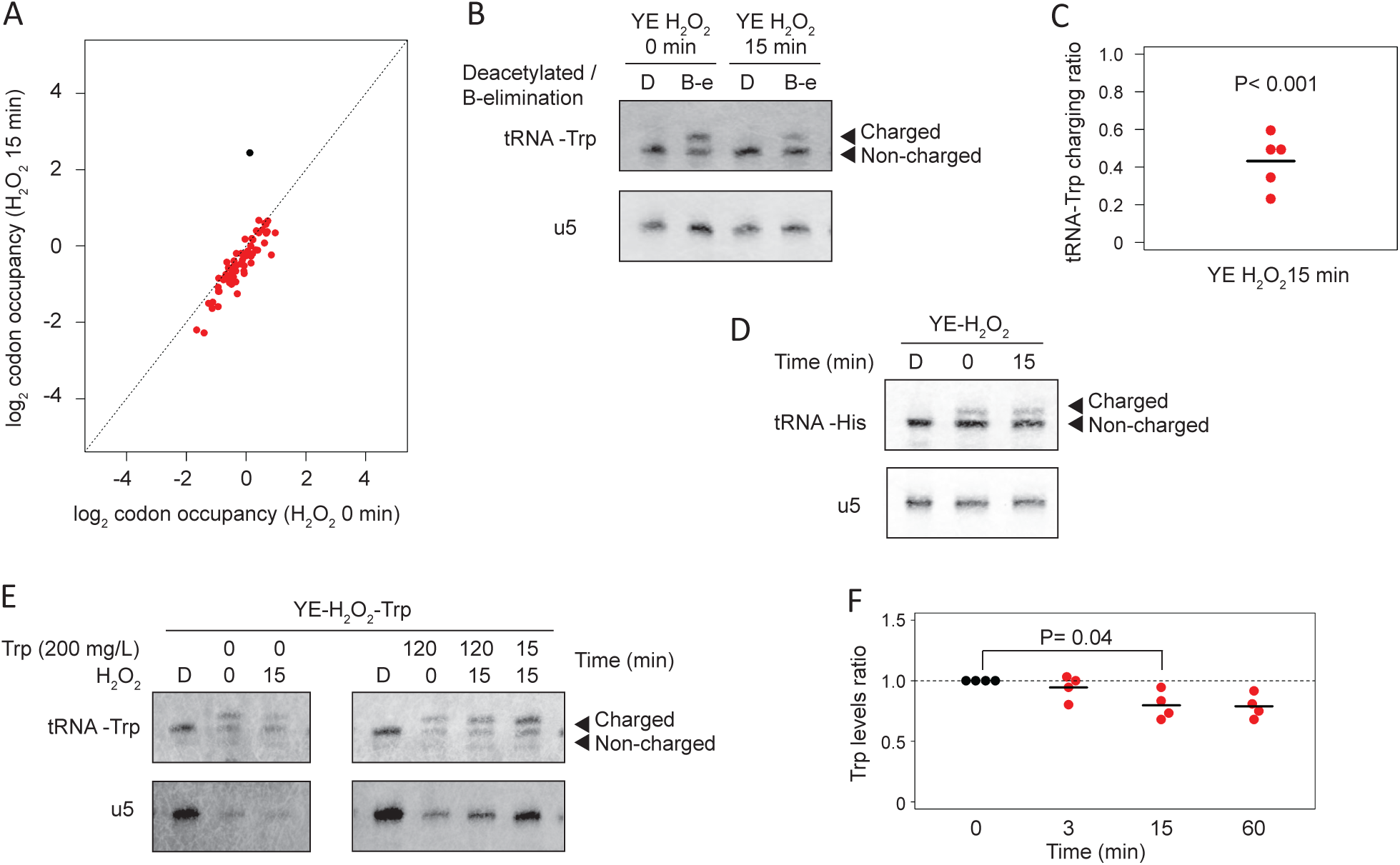
Levels of charged tRNA-Trp are affected by oxidative stress. **(A)** Scatter plots showing log_2_ relative codon enrichments before and after H_2_O_2_ treatment for 15 minutes in wild type cells. The TGG codon encoding tryptophan is plotted in black. **(B)** Representative northern blot for the determination of tRNA-Trp charging levels before and after H_2_O_2_ exposure. The top blot was hybridised with a probe against tRNA-Trp, and the bottom one with a probe against the U5 snRNA. In the upper blot, the top band corresponds to charged tRNA, and the bottom to the uncharged form. tRNA-Trp samples were either deacylated to remove the linked amino acid from charged tRNAs (sample D) or oxidised to remove the unprotected 3’ nucleotides from uncharged tRNAs by beta-elimination (sample B-e) (see Methods for details). U5 snRNA was used as a loading control. **(C)** Quantification of tRNA-Trp charging ratios. Ratios between charged and uncharged tRNA were calculated, and normalised to the ratio in untreated cells. Each dot corresponds to an independent biological replicate (n = 5), and the horizontal line indicates the mean. Significance was calculated by using a paired Student’s t test. **(D)** As in C, but using a probe against tRNA-His (top panel) or U5 snRNA (bottom). **(E)** Northern blot as in C, to explore the effects of supplementing the culture medium with tryptophan. Cells were grown in the presence of tryptophan for 0, 15 or 120 min, and H_2_O_2_ was added at the indicated times (0, 15 min) before the end of the incubation with tryptophan. Control deacylated RNA (sample D) is used to identify the location of uncharged tRNA. **(F)** Changes in intracellular tryptophan levels in response to H_2_O_2_ exposure. Tryptophan levels were measured at the indicated times after H_2_O_2_ addition to the culture medium. Each dot corresponds to an independent biological replicate (n = 4), and the horizontal lines indicate the means. Significance was calculated using a paired Student’s t test. No adjustment for multiple testing was performed.

A possible explanation for this observation was that cellular tryptophan levels decreased because of H_2_O_2_ treatment. We therefore measured intracellular amino acid concentrations by mass spectrometry. We observed a reduction of approximately 20% in tryptophan levels. However, this change was borderline of statistically significance (P=0.04), and was smaller than that of other amino acids that did not show any difference in ribosome occupancy of their cognate codons (Fig. 5F, S4C). In addition, mRNA levels and TE of the tRNA-Trp ligase, encoded by *wrs1* gene, remained unaffected.

A second hypothesis was that the levels of tRNA charging could be affected by oxidative stress. To investigate this possibility, we compared the levels of amino-acylated (charged) and deacylated (uncharged) tRNA-Trp. tRNAs were first subjected to periodate oxidation. This treatment leads to the removal of the 3’ nucleotide of uncharged tRNAs through β-elimination, whereas the charged fraction is protected by the amino acid and remains unaltered. The tRNAs are then deacylated at high pH. Thus, uncharged and charged tRNAs show a difference of one nucleotide in size, which can be detected by polyacrylamide gel electrophoresis and Northern blotting [51] (Fig. 5B). tRNA-Trp deacylated samples were used as controls and showed that total levels of tRNA-Trp were not affected by oxidative stress, as well as ruling out the fragmentation of tRNAs [27] (Fig. 5B). By contrast, tRNA-Trp charging levels were reduced more than two-fold after H_2_O_2_ treatment (Fig. 5B, C). To confirm that this effect is specific of tRNA-Trp, we verified that charged tRNA-His levels remained identical upon H_2_O_2_ treatment (Fig. 5D). The addition of supplemental tryptophan to rich medium two hours before, or only during the H_2_O_2_ treatment, prevented the charged tRNA-Trp drop and increased the charging levels (Fig. 5E). Thus, these data indicate that the increase in ribosome occupancy at the tryptophan codon in H_2_O_2_ stress condition reflects a reduced charged tRNA-Trp fraction.

### Decreased tRNA-Trp charging may affect eIF2α phosphorylation

In *S. cerevisiae* uncharged tRNAs activate the Gcn2 kinase, which phosphorylates eIF2α to downregulate global translation initiation in response to amino acid starvation [7]. Recently, it has been shown that Gcn2 in *S. pombe* is activated in response to UV and oxidative stress through a mechanism that involves Gcn1 and most likely the binding of tRNAs [52]. Thus, we reasoned that increased levels of uncharged tRNA-Trp upon H_2_O_2_ treatment might contribute to eIF2α phosphorylation. To explore this possibility, we compared eIF2α phosphorylation levels upon H_2_O_2_ stress in the presence and absence of supplemental tryptophan in rich medium. Consistent with this idea, addition of tryptophan (which increases tRNA-Trp charging levels, see above) caused a reduction of eIF2α phosphorylation (Fig. 6C). These data suggest that uncharged tRNA-Trp could promote Integrated Stress Response signalling upon H_2_O_2_ exposure in rich medium, and thus affect both translation initiation and translation elongation. Differences in charged tRNA-Trp levels cannot affect codon usage because tryptophan is encoded by only one codon, TGG. However, as tryptophan is the least frequent amino acid in proteins (less than 0.015% on average), we asked whether the translation efficiency of genes containing more tryptophan might be affected by lower charged tRNA-Trp levels upon H_2_O_2_ stress. Genes were ranked by the percentage of tryptophan codons in their coding sequences, and assigned to 11 bins. We then measured the apparent change in translation efficiency upon oxidative stress for each bin. The data showed a trend towards increased translation efficiency changes after H_2_O_2_ treatment with higher tryptophan content, which was not observed in other stresses (Fig. 6A and Fig. S5). We also ruled out that this trend was determined by transcript lengths (Fig. 6B). The group of genes lacking tryptophan is enriched in genes encoding ribosomal proteins (P = 10^−27^), which have a small transcript size (Fig. 6B).

**Fig. 6.**
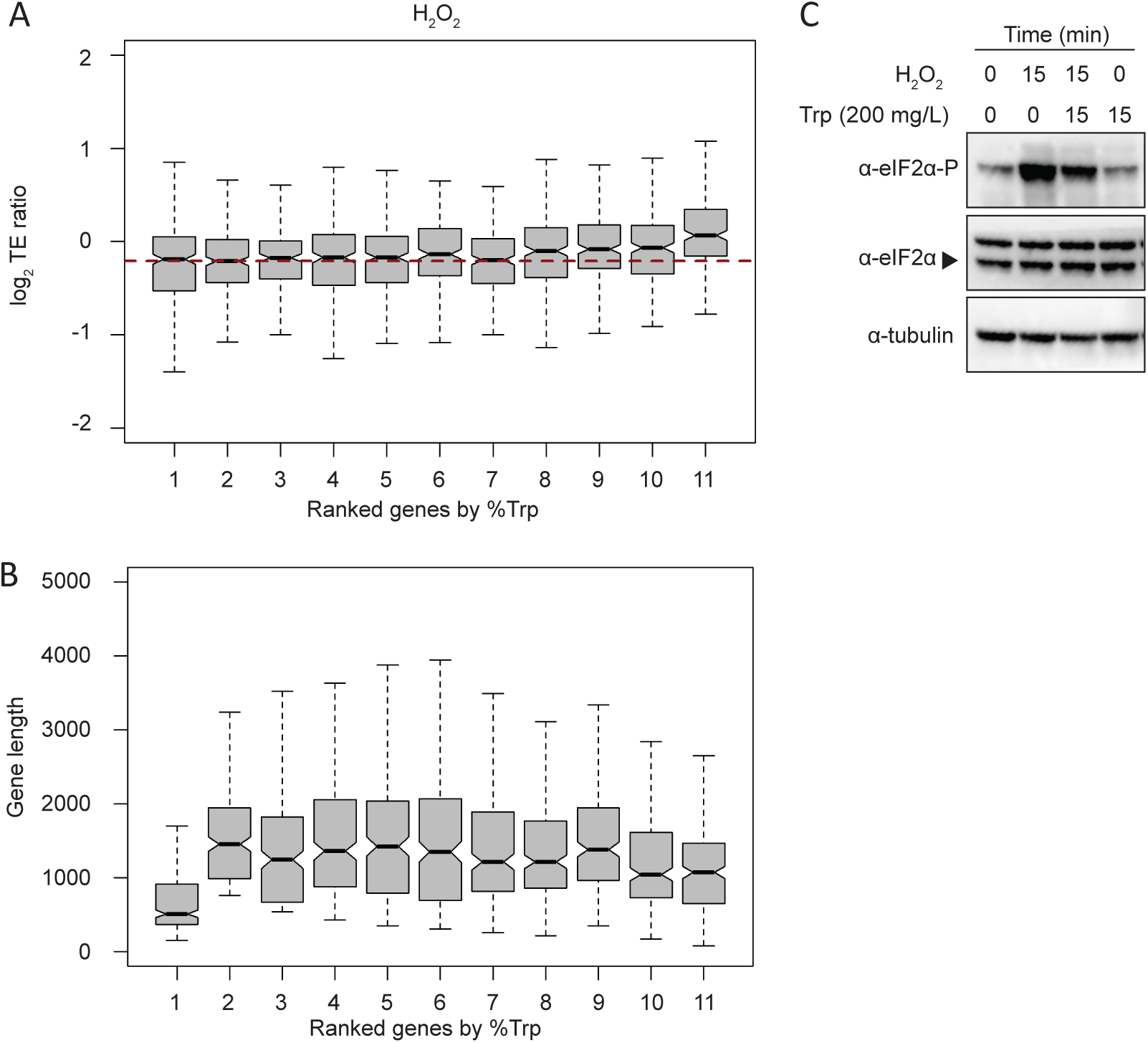
Oxidative stress affects the translation efficiency of tryptophan-enriched genes, and decreased tRNA-Trp charging may affect eIF2α phosphorylation. **(A)** Boxplots showing changes in translation efficiency upon oxidative stress (log_2_ TE ratios stress/control, 15 min treatment) according to tryptophan content. Genes were binned into 11 categories based on the fraction of tryptophan in their coding sequences (the first group contains 269 genes without tryptophan, and the other 10 groups have 234 genes each). The horizontal red dashed line indicates the median of the second group. **(B)** As above, but displaying coding sequence lengths. **(C)** Western blots to investigate the effect of tryptophan on eIF2α phosphorylation levels after H_2_O_2_ treatment. Cells were treated with H_2_O_2_ for 15 minutes and supplemental tryptophan was added as indicated. Tubulin was used as a loading control.

By contrast, the group with the most tryptophan (0.023-0.056%) showed an enrichment in genes related to lipid biosynthesis (GO:0008610; P = 6.6× 10^−10^), protein glycosylation (GO:0006486; P = 4× 10^−7^) and cell wall biogenesis (GO:0071554; P = 1.5× 10^−5^). This enrichment is consistent with the fact that tryptophan is an amphipathic amino acid, often found in transmembrane domains. Given that translation efficiency as defined above is a relative measurement of the number of ribosomes per transcript (normalised to mRNA abundance), an increase in this parameter in tryptophan-rich genes is likely to reflect a slowdown of translation elongation of these genes rather than an increase in their protein synthesis. Thus, we propose that oxidative stress modulates translation at two levels: at initiation, through the Integrated Stress Response, and at elongation, through tryptophan stalling.

## DISCUSSION

### The translational landscape of the response to multiple stresses

We present a genome-wide analysis of the translational response of the fission yeast *S. pombe* to five different stresses using highly standardized and comparable conditions. We found that heavy metal exposure, heat shock and oxidative stress led to stronger transcriptional and translational changes than DNA damage and osmotic stress. Moreover, increased levels of eIF2α phosphorylation correlate with pronounced global downregulation of translation and with strong induction of *fil1* translation. Of course, as we monitored a single time point and condition for each stress, the conclusions about the relative strength of the effects are only valid to the specific experimental conditions used. Despite this caveat, our results indicate that the translational response to stress is mainly directed to repress the translational machinery, with few genes upregulated at this level. Moreover, we found that many differentially translated genes are often regulated in multiple stresses. Thus, our study identifies the key players of a stress response at the translational level and provides insight into the biological response to stress.

### Fil1 regulation and role in stress responses

The induction of the Fil1 transcription factor in multiple stress conditions in rich medium was surprising, as Fil1 is a master regulator of the amino acid starvation response (analogous to Gcn4 and Atf4), and is required to maintain a normal growth rate in minimal medium [12]. Cells lacking Fil1 showed similar growth than wild type cells in rich medium, suggesting that Fil1 does not have a role in unstressed cells in rich medium. Strikingly, the *fil1* gene showed much higher translational induction upon cadmium, heat shock and H_2_O_2_ in rich medium than in amino acid starvation induced by 3-AT in minimal medium (17-fold, 12-fold, 13-fold and 3.8-fold, respectively). A possible explanation is that *fil1* translation levels may already be higher in minimal medium in unstressed cells. Similarly, the induction of the Fil1 orthologue in *S. cerevisiae* (Gcn4) has been reported not only in amino acid starvation or glucose limitation, but also after MMS (although the conditions were different from the ones used in this work) and H_2_O_2_ treatments [33,34,53]. These results suggest that metabolic adaptation, mediated by translationally controlled transcription factors such as Fil1 and Gcn4, is an evolutionary conserved part of many stress responses (and not just amino acid starvation).

Fil1-dependent genes are mostly related to amino acid metabolism and transmembrane transport, and show a significant overlap with CESR induced genes. In rich medium, cells lacking Fil1 are unable to regulate the expression of Fil1 targets after cadmium, heat shock and H_2_O_2_ treatments. In addition, this regulation is also impaired in *eIF2α-S52A* cells, consistent with the complete absence of *fil1* induction. Surprisingly, Fil1 and its targets were upregulated in *sty1Δ* cells (compared to wild type) in unstressed conditions, possibly because the lack of a normal transcriptional response in these cells may lead to stress [14]. Moreover, it has been described that *sty1Δ* cells are sensitive to stress [45–48]. This may also reflect that Sty1 may have a role in the modulation of eIF2α kinases after stress. Indeed, *sty1Δ* cells show increased eIF2α phosphorylation after oxidative stress, which might result in induction of Fil1 [10,21]. Given that Sty1 is also important in adaptation to stress and that nothing is known about Fil1 protein stability or post-translational modifications, it could be also involved in downregulation of Fil1 after stress.

Strikingly, upon cadmium treatment, despite the induction of Fil1, the expression of Fil1-dependent genes in wild-type cells is very weak and *fil1Δ* cells are resistant to this stress. Cadmium stress increase ROS and intracellular oxidative stress [42], but the response is very different to H_2_O_2_ treatment. For example, cadmium is imported through specific transporters, whereas H_2_O_2_ freely diffuses into cells. Moreover, H_2_O_2_ converts into superoxide, and is a substrate for the peroxiredoxin system that is responsible for the oxidation of most proteins. Indeed, *S. pombe* cells lacking transcription factors like Atf1 or Prr1 are sensitive to H_2_O_2_, but resistant or insensitive to cadmium [39,54]. Additionally, it was reported that Fil1 might drive the response to amino acid starvation partially through the action of downstream transcription factors [12]. Thus, different kinetics in the induction and the expression of Fil1-dependent genes suggest that Fil1 might be modulating transcriptional changes depending on the stress through other transcription factors. We propose that Fil1 acts as a master regulator of several stress conditions, promoting a distinct response for each situation. Further work will be required to unveil the direct targets of Fil1 under each stress condition, and whether Fil1 can activated them in different ways to modulate specific responses.

### Ribosomes stall on tryptophan codons upon H_2_O_2_ treatment

Ribo-seq experiments revealed that oxidative stress caused ribosome stalling on tryptophan codons, which correlated with decreased levels of charged tRNA-Trp. By contrast, intracellular levels of tryptophan were not changed significantly under these conditions, suggesting that tryptophan metabolism is not directly affected by the stress. This suggests that changes in tRNA modifications, which are poorly described in *S. pombe*, or in the tRNA-Trp synthetase activity, may affect the levels of charged tRNA-Trp and cause stalling.

The main eIF2α kinase activated in response to early exposure to H_2_O_2_ in *S. pombe* is Gcn2 [10]. The molecular pathway leading to activation of Gcn2 upon nutrition limitation is well understood: Gcn2 is activated by uncharged tRNAs, which bind the histidyl-tRNA synthetase-related domain of Gcn2 at the C-terminus [55–57]. However, Gcn2 is activated by other stresses that are not expected to cause uncharged tRNA accumulation [52], such as UV irradiation or oxidative stress. Under these conditions, the tRNA binding domain of Gcn2 is still required for its activation [52]. Here, we provide evidence for the first time that a nutrient-unrelated stress like oxidative stress causes uncharged tRNA accumulation. In addition, we show that eIF2α phosphorylation after oxidative stress correlates with uncharged tRNA-Trp accumulation; addition of tryptophan results in a reduction in the levels of both uncharged tRNA-Trp and of eIF2α phosphorylation. Thus, we propose that the accumulation of uncharged tRNA-Trp upon oxidative stress, which causes a mild elongation defect, is used to regulate initiation through Gcn2 activation.

## METHODS

### Strains, Growth Conditions, and Experimental Design

All strains used were prototrophic. Table S1 presents a full list of strains. Standard methods and media were used for *S. pombe* [58]. For all genome-wide stress experiments, *S. pombe* cells were grown in YES medium (supplemented with leucine, uracil and adenine) at 32 °C. Cells were treated for 15 minutes as described below. Heavy metal stress: cadmium sulphate (CdSO_4_; 481882; Sigma) was added to a final concentration of 0.5 mM. Oxidative stress: hydrogen peroxide (H_2_O_2_; H1009; Sigma) was added to a final concentration 0.5 mM. Heat stress: cells were quickly transferred from 32°C to a prewarmed flask in a 39°C water bath. Alkylating agent: methyl methanesulfonate (MMS, 129925, Sigma) at a final concentration of 0.02%. Osmotic stress: cells were grown to OD_600_ = 0.7, and diluted with prewarmed YES 3 M sorbitol (BP-439-500, Fisher) to a final concentration 1 M sorbitol.

For plate drop assays, cells were grown in YES to exponential phase at 32 °C and plated in 10-fold dilutions. Plates were incubated for two days at 32 °C (except the plate incubated at 39 °C for heat shock assays). For tryptophan charging experiments, supplemental tryptophan (DOC0188, Formedium) was added to the culture from a stock 8 g/L in YES to a final concentration of 200 mg/L.

All repeats of genome-wide experiments were independent biological replicates carried out on separate days (see ArrayExpress deposition footnote for a complete list). The following sequencing experiments were performed: (i) ribosome profiling and matching RNA-seq in five stress conditions of three strains (wild-type, *eIF2α-S52A* and *sty1Δ*), and (ii) RNA-seq of *fil1Δ* and cells in five stress conditions.

### Amino acid analysis

Amino acid quantification was performed by liquid chromatography selective reaction monitoring (LC-SRM) as described [59].

### Protein Analyses

To prepare samples for Western blotting, cells were harvested by filtration, washed with 20% TCA, resuspended in 100 μL of 20% TCA, and frozen. Cell pellets were lysed with 1 mL of acid-treated glass beads in a bead beater (FastPrep-5; MP Biomedicals) at level 7.5 for 15 s, and 400 μL of 5% TCA was added before eluting from the glass beads. Lysates were frozen on dry ice and spun at 18,000 relative centrifugal force for 10 min. Pellets were resuspended in SDS-Tris solution (2% SDS and Tris 0.3M pH 10.7), boiled during 5 min and cleared by centrifugation at maximum speed for 2 min. Protein extract concentrations were measured using Pierce BCA Protein Assay solution (Thermo), and 60-90 µg of total protein were loaded with Laemmli SDS Sample Buffer (reducing) (Alfa Aesar). For western blot analysis, the following antibodies were used: anti-eIF2α (1:500; 9722, Cell Signalling), anti-Phospho-eIF2α (1:1,000; 9721, Cell Signalling), and anti-tubulin (1:10,000; sc-23948, Santa Cruz). The secondary antibodies were HRP-conjugated goat anti-mouse IgG (H+L) (1:10,000; 31430, Thermo Fisher) and anti-rabbit IgG (1:10,000; ab-6721, Abcam). TAP tag was detected with peroxide–anti-peroxide complexes (P1291; Sigma). Detection was performed using the enhanced chemiluminescence procedure (ECL kit).

### Northern analysis of aminoacyl-tRNA charging

RNA from the indicated conditions was prepared by phenol extraction under acidic conditions using acid buffer (0.3 M sodium acetate pH 5, 10 mM Na_2_EDTA) for resuspension of frozen cells, and final acid RNA buffer (10 mM sodium acetate pH 5, 1mM EDTA). As control samples, RNA was deacylated with 0.2 M Tris pH 9.0 for 2h at 37 °C. Periodate oxidation and β-elimination were performed as described [51] using 2 µg of total RNA. In the final step, all samples were deacylated. tRNAs were detected by Northern blotting in non-acidic conditions: RNA samples were loaded onto a 6.5 % denaturing urea polyacrylamide gel, electrophoresed at 17 W for 70 min and transferred onto an Amersham Hybond-N membrane (GE Healthcare) using the Trans-blot SD semi-dry transfer cell (Biorad) as described [60], but using non-acidic conditions. The oligonucleotide probes used were: tRNA Trp CCA (5’-TGACCCCTAAGTGACTTGAACACTTGA-3’), tRNA His GUG (5’-TGCCCACACCAGGAATCGAACCTGGGT-3’) and U5 snRNA as loading control (5’-GCACACCTTACAAACGGCTGTTTCTG-3’) [51]. The oligonucleotides were labelled with infrared dyes (IRD-700 or IRD-800) at the 5’ end for fluorescence detection using the LICOR Odyssey system. The tRNA loading was quantified as the ratio of upper to lower bands relative to the unstressed condition.

### Polysome profiling, ribosome profiling, library preparation and sequencing

Ribosome-protected fragment (RPF) analyses, preparation of cell extracts, RNase treatment, separation of samples by centrifugation through sucrose gradients, and isolation of protected RNA fragments were performed as described [61]. For polysome profiles, there is no RNase I digestion step, and lysis buffer and sucrose solutions were prepared with double concentration of MgCl_2_ (10 mM). Polysome-subpolysome ratio was quantified by measuring the area under the curve relative to the unstressed conditions using Image J software (NIH).

For all RPF samples, gel-purified RNA fragments of around 17-30 nucleotides were treated with 10 units of T4 PNK (Thermo Fisher) in a low-pH buffer (700 mM Tris, pH 7, 50 mM DTT, and 100 mM MgCl_2_) for 30 min at 37 °C. ATP and buffer A (Thermo Fisher) were then added for an additional 30 min incubation. RNA fragments were column-purified (PureLink RNA microcolumns; Life Technologies). A total of 100 ng was used as input for the NEXTFLEX Small RNA Sequencing Kit (Version 3; Bioo Scientific), and libraries were generated following the manufacturer’s protocol. For mRNA analyses, total RNA was isolated as described [61]. Total RNA was then depleted from rRNA by using Ribo-Zero Gold rRNA Removal Kit Yeast (Illumina) with 4 μg as input. Finally, 30 ng of rRNA-depleted RNA was used as starting material for the NEXTflex Rapid Directional qRNA-Seq Kit (Bioo Scientific). Libraries were sequenced in an Illumina HiSeq4000 or Novaseq6000 as indicated (ArrayExpress submission).

### Data Analysis

Data processing and read alignment were performed as described [12]. Data quantification (number of reads per coding sequence) was carried out by using in-house Perl scripts as described [12]. All statistical analyses were performed using R.

Differential expression analysis was performed by using the Bioconductor DESeq2 package [62]. Raw counts were directly fed to the program, and no filtering was applied. Unless otherwise indicated, a threshold of 10^−2^ was chosen for the adjusted P value, and a cut-off of 2-fold minimal change in RNA levels.

For codon usage analyses, RPF reads were aligned to nucleotide 13 (corresponding to position 1 of the codon in the ribosome P site). Only codons after 90 were used. For each coding sequence, the following calculations were performed: (i) determination of the fraction of RPFs that occupy each codon (RPFs in a given codon divided by total RPFs); (ii) quantification of the relative abundance of each codon on the coding sequence (number of times each codon is present divided by total codon number); and (iii) definition of the normalized codon occupancy by dividing parameter 1 by parameter 2. The average codon enrichments (Fig. 5, S4) were then calculated with data from all coding sequences.

For the analysis of translational efficiencies we used RiboDiff [63]. RiboDiff was provided with raw read counts for each gene, from ribosome profiling and from RNA-seq. To select differentially translated genes, a threshold of 10^−2^ for the adjusted P value, and a cut-off of 1.5-fold were chosen.

Translation efficiency and mRNA ratios were median-centred for plotting. The list of Fil1 targets for Figures 4 and S3 was obtained from, Dataset_S01, repressed genes in *fil1Δ versus* wild type without stress (no 3AT) [12] and only those genes from the lists with at least 20 counts in 80% or more samples were used. Gene set enrichment was performed with AnGeLi [64]. The significance of the overlap between gene lists was calculated using Fisher’s exact test.

### Data availability

All raw data files have been deposited in ArrayExpress [65] [https://www.ebi.ac.uk/arrayexpress/] under accessions: E-MTAB-8746, E-MTAB-8686, E-MTAB-8744, E-MTAB-8745, E-MTAB-8602 and E-MTAB-8583.

## Supporting information

Dataset_S1_Induced_and_repressed_gene_expression

Dataset_S2_Induced_and_repressed_TE

Dataset_S3_mRNA_counts

Dataset_S4_RPF_counts

## ACKNOWLEDGMENTS

This work was supported by a Biotechnology and Biological Sciences (BBSRC) grant to Juan Mata (BB/N007697/1). Markus Ralser was supported by the Francis Crick Institute, which receives its core funding from Cancer Research UK (FC001134), the UK Medical Research Council (FC001134), and the Wellcome Trust (FC001134), as well as specific project funding from the Wellcome Trust (IA 200829/Z/16/Z to M.R.), as well as the German Federal Ministry of Education and Research (BMBF) as part of the National Research Node Mass Spectrometry in Systems Medicine (MSCoreSys 031L0220A). We thank Samuel Marguerat and Caia Duncan for comments on the manuscript and Mathew Peacey and Anno Koetje for help with experiments.

## AUTOR CONTRIBUTIONS

A.R., S.G. and J.M. designed the study. A.R., S.G. and M.M. performed and analysed experiments. A.R., M.R. and J.M. analysed data. A.R. and J.M. wrote the manuscript.

---

**Table S1:**
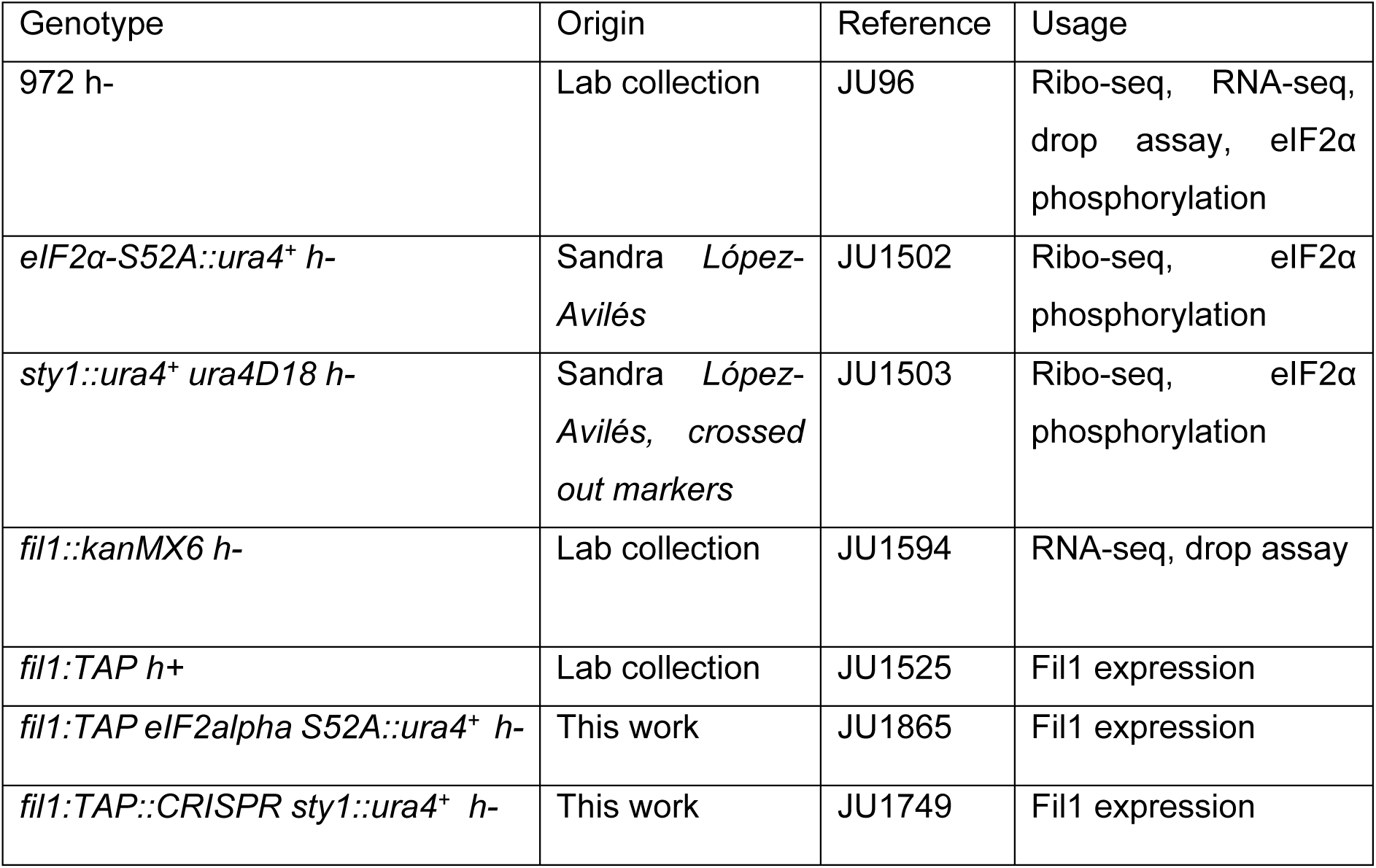
Strain and experiment list.

| Genotype | Origin | Reference | Usage |
| --- | --- | --- | --- |
| 972 h- | Lab collection | JU96 | Ribo-seq, RNA-seq, drop assay, eIF2α phosphorylation |
| *eIF2α-S52A::ura4<sup>+</sup> h-* | Sandra López-Avilés | JU1502 | Ribo-seq, eIF2α phosphorylation |
| *sty1::ura4<sup>+</sup> ura4D18 h-* | Sandra López-Avilés, crossed out markers | JU1503 | Ribo-seq, eIF2α phosphorylation |
| *fil1::kanMX6 h-* | Lab collection | JU1594 | RNA-seq, drop assay |
| *fil1:TAP h+* | Lab collection | JU1525 | Fil1 expression |
| *fil1:TAP eIF2α S52A::ura4<sup>+</sup> h-* | This work | JU1865 | Fil1 expression |
| *fil1:TAP::CRISPR sty1::ura4<sup>+</sup> h-* | This work | JU1749 | Fil1 expression |

**Fig. S1.**
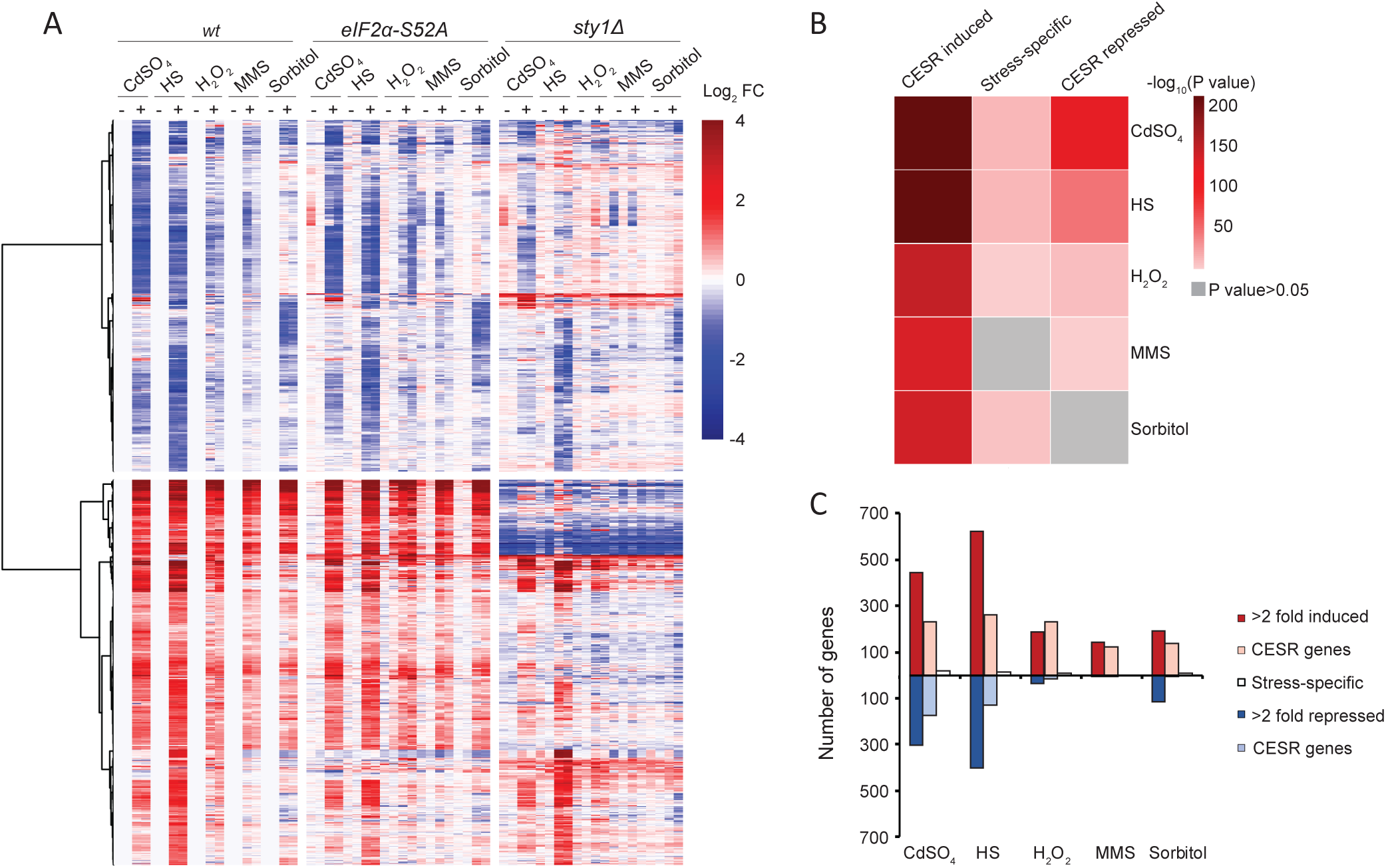
Transcriptomic responses to stress. **(A)** Heat map of changes in mRNA levels in response to five stress conditions (log_2_ ratio stress/control). Data are shown for 1,247 genes that are differentially expressed in at least one stress in wild type (see Methods for details). All data are normalised to the corresponding untreated wild type sample. **(B)** Heat map displaying enrichment analysis of previously described CESR gene lists [14] and genes differentially expressed during five stress treatments (our dataset). **(C)** Comparison of the absolute numbers of induced and repressed genes in our experiments (>2-fold repressed or induced) with the CESR-induced, CESR-repressed and stress-specific genes [14].

**Fig. S2.**
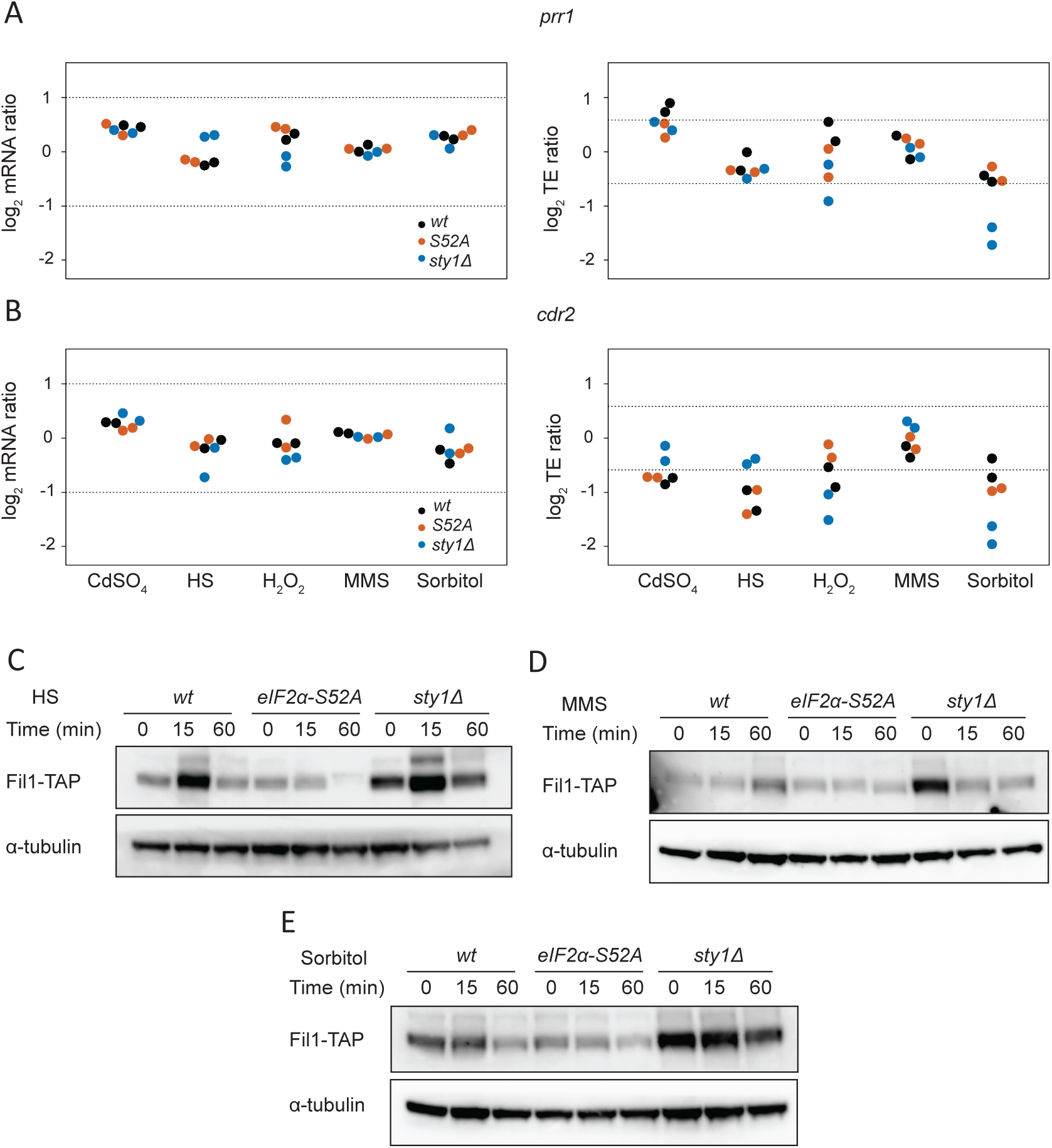
Gene-specific translational regulation. **(A)** Changes in translation efficiency and transcript levels (log_2_ ratios stress/control) obtained for the *prr1* gene. Data for two biological replicates are shown. **(B)** As in A, but for the *cdr2* gene. **(C to E)** Western blots to measure Fil1-TAP protein levels after heat shock, MMS and sorbitol at the indicated times and genetic backgrounds. Tubulin was used as a loading control.

**Fig. S3.**
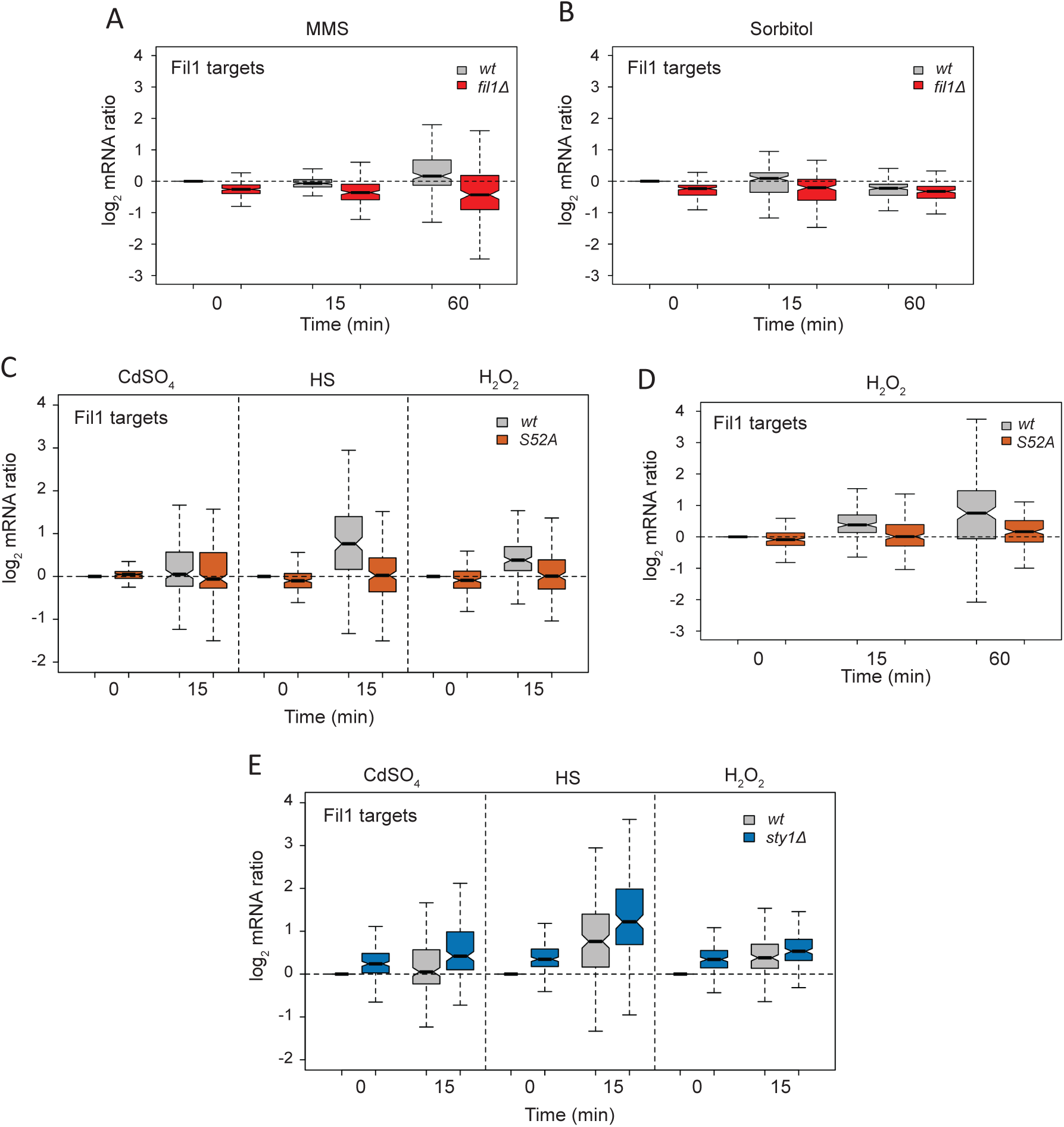
Transcription of Fil1 targets relies on eIF2α phosphorylation and is increased in unstressed *sty1Δ* cells. **(A and B)** Boxplot showing changes in mRNA levels (log_2_ ratio stress/control) of genes encoding Fil1 targets after MMS and sorbitol treatments, in wild type and *fil1Δ* strains. **(C)** As in A and B, but after cadmium, heat shock and H_2_O_2_ treatments, in wild type and *eIF2α-S52A* strains. **(D)** As in C, but after H_2_O_2_ treatment and with an additional time point. **(E)** As in C, but in the wild type and *sty1Δ* strains.

**Fig. S4.**
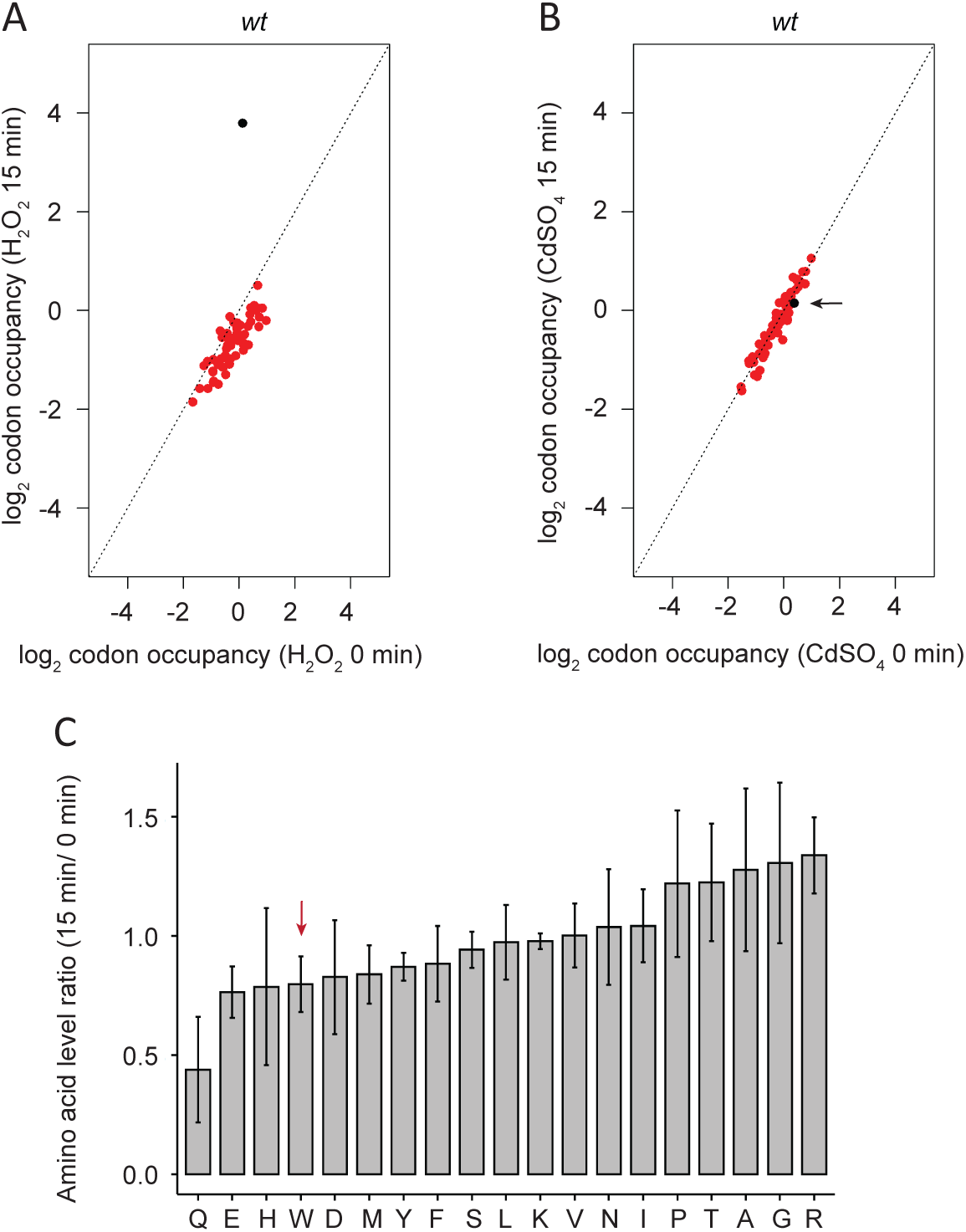
Tryptophan codon enrichment is specific for H_2_O_2_ treatment. **(A)** Scatter plots showing log_2_ relative codon enrichments before and after H_2_O_2_ treatment for 15 minutes in wild type cells (similar to Fig. 5A, but a different biological replicate). The TGG codon encoding tryptophan is plotted in black. **(B)** As in A, after cadmium treatment. TGG is displayed in black and highlighted with an arrow. **(C)** Change in amino acid levels (ratio stress/control) upon oxidative stress exposure. Data are from 4 independent biological replicates (means ± SD). Tryptophan (W) is highlighted by a red arrow.

**Fig. S5.**
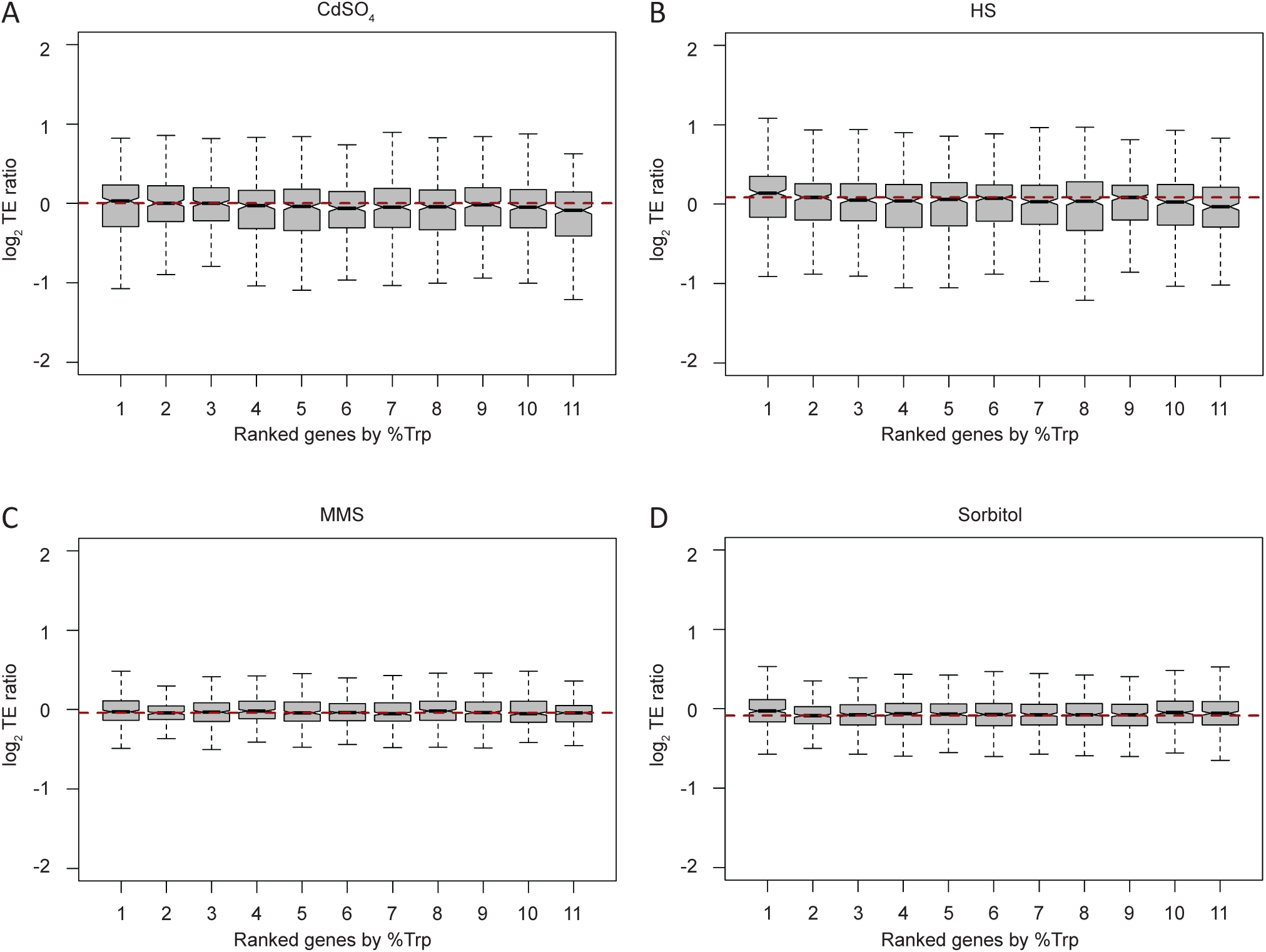
Translation efficiency of tryptophan-enriched genes after other stress treatments. **(A to D)** Boxplots displaying changes in translation efficiency upon the indicated stresses (log_2_ stress/control ratios) according to tryptophan content. Genes were binned into 11 categories based on the fraction of tryptophan in their coding sequences (the first group contains 269 genes lacking tryptophan, and the other 10 groups have 234 genes each). The horizontal red dashed line indicates the median of the second group.

